# Snapshot of 5-HT_2A_ receptor activation in the mouse brain via IP_1_ detection

**DOI:** 10.1101/2024.10.11.617861

**Authors:** Mario de la Fuente Revenga, Javier González-Maeso

## Abstract

The distinct subjective effects that define psychedelics such as LSD, psilocybin or DOI as drug class are causally linked to activation of the serotonin 2A receptor (5-HT_2A_R). However, some aspects of 5-HT_2A_R pharmacology remain elusive, such as what molecular drivers differentiate psychedelic from non-psychedelic 5-HT_2A_R agonists. We developed an ex vivo platform to obtain snapshots of drug-mediated 5-HT_2A_R engagement of the canonical G_q/11_ pathway in native tissue. This non-radioactive methodology captures the pharmacokinetic and pharmacodynamic events leading up to changes in inositol monophosphate (IP_1_) in the mouse brain. The specificity of this method was assessed by comparing IP_1_ levels in homogenates from the frontal cortex in DOI-treated wild-type and 5-HT_2A_R-KO animals compared to other brain regions, namely striatum and cerebellum. Furthermore, we encountered that head-twitch response (HTR) counts and IP_1_ in the frontal cortex were correlated. We observed that IP_1_ levels in frontal cortex homogenates from mice treated with LSD and lisuride vary in magnitude, consistent with LSD’s 5-HT_2A_R agonism and psychedelic nature, and lisuride’s lack thereof. MDMA evoked an increase of IP_1_ signal in the frontal cortex that were not matched by the serotonin precursor 5-HTP or the serotonin reuptake inhibitor fluoxetine. We attribute differences in the readout primarily to the indirect stimulation of 5-HT_2A_R by MDMA via serotonin release from its presynaptic terminals. This methodology enables capturing a snapshot of IP_1_ turnover in the mouse brain that can provide mechanistic insights in the study of psychedelics and other serotonergic agents pharmacodynamics.

## Introduction

The last decade has witnessed a burgeoning renewed interest in psychedelic drugs. Neglected for decades outside the underground and counter-cultural movements, drugs such as lysergic acid diethylamide (LSD) *N,N*-dimethyltryptamine (DMT), psilocybin, mescaline, and other synthetic derivatives reappear as a research priority as the result of preliminary evidence suggesting that they may serve as treatments in mental health diagnoses for which therapeutic options are limited^1^. Albeit subjective in nature, the characteristic altered mental state elicited by psychedelics is attributable to discrete molecular interactions with specific neuronal receptors^2^. Mounting evidence from studies in humans, animal models and in vitro assays demonstrate the central role of serotonin (or 5-hydroxytryptamine) 2A (5-HT_2A_R) receptor activation in the action of classical psychedelics^3–6^.

5-HT2AR is a G protein-coupled receptor (GPCR) that canonically engages upon activation G_q/11_ proteins primarily^6^. Downstream, recruitment of these mainly excitatory heterotrimeric G proteins results in the production of inositol triphosphate (IP_3_), with subsequent release of calcium from intracellular compartments. This mechanism is consistent with the hyperexcitable state that neurons enter when exposed to psychedelics such as LSD, and with the apparent increase in cortical metabolic activity associated with psilocybin administration in healthy human volunteers^7–9^. However, 5-HT_2A_R activation is not exclusively attained by psychedelics. Lisuride, 2-Br-LSD, Ariadne and 6-F-DET are a small but representative sample of compounds known to activate 2A but lack in the subjective effects of psychedelics^10–13^

There is an ongoing need for functional benchmarks that can link psychedelics and their manifestations to their underlying molecular actions. Connecting these domains can aid to identify potential therapeutics pathways and provide much clarity on the mechanistic drivers that differentiate psychedelic from non- or lesser psychedelic 5-HT_2A_R agonists^14,15^. With this goal in mind, we adapted a commercial a methodology based on homogenous-time resolved fluorescence (HTRF) for the determination of inositol phosphate (IP_1_)—a downstream metabolite of G_q_ and related G protein-dependent signaling pathways—from mouse brain dissects. The mouse brain samples are subject to minimal post-extraction manipulation involving only homogenization. In doing so, we managed to obtain a molecular readout from the mouse brain that, like in vitro systems, is representative of 5-HT_2A_R activation by psychedelics. But unlike in in vitro methods is also subject to the pharmacokinetics that determine drug action in vivo.

## Results

### IP_1_ detection in mouse brain samples

IP_1_ is a near-terminal metabolite product resulting from the sequential hydrolysis of IP_3_; a key second messenger molecule in the signal transduction of receptors like 5-HT_2A_R that couple to G_q/10/11/14/15_ (G_q_ for brevity)^16^. HTRF detection of IP_1_ can readily be employed as a reporter of G_q_ activity in vitro in cultured cells using commercially available kits. Compelled by previous attempts to study IP_1_ production by muscarinic receptors in vivo^17^, we aimed to adapt detection of IP1 via HTRF to detect changes in levels of this metabolite in mouse brain homogenates from small samples from 5-HT_2A_R-rich brain regions(**Fig 1**).

**Figure 1.**
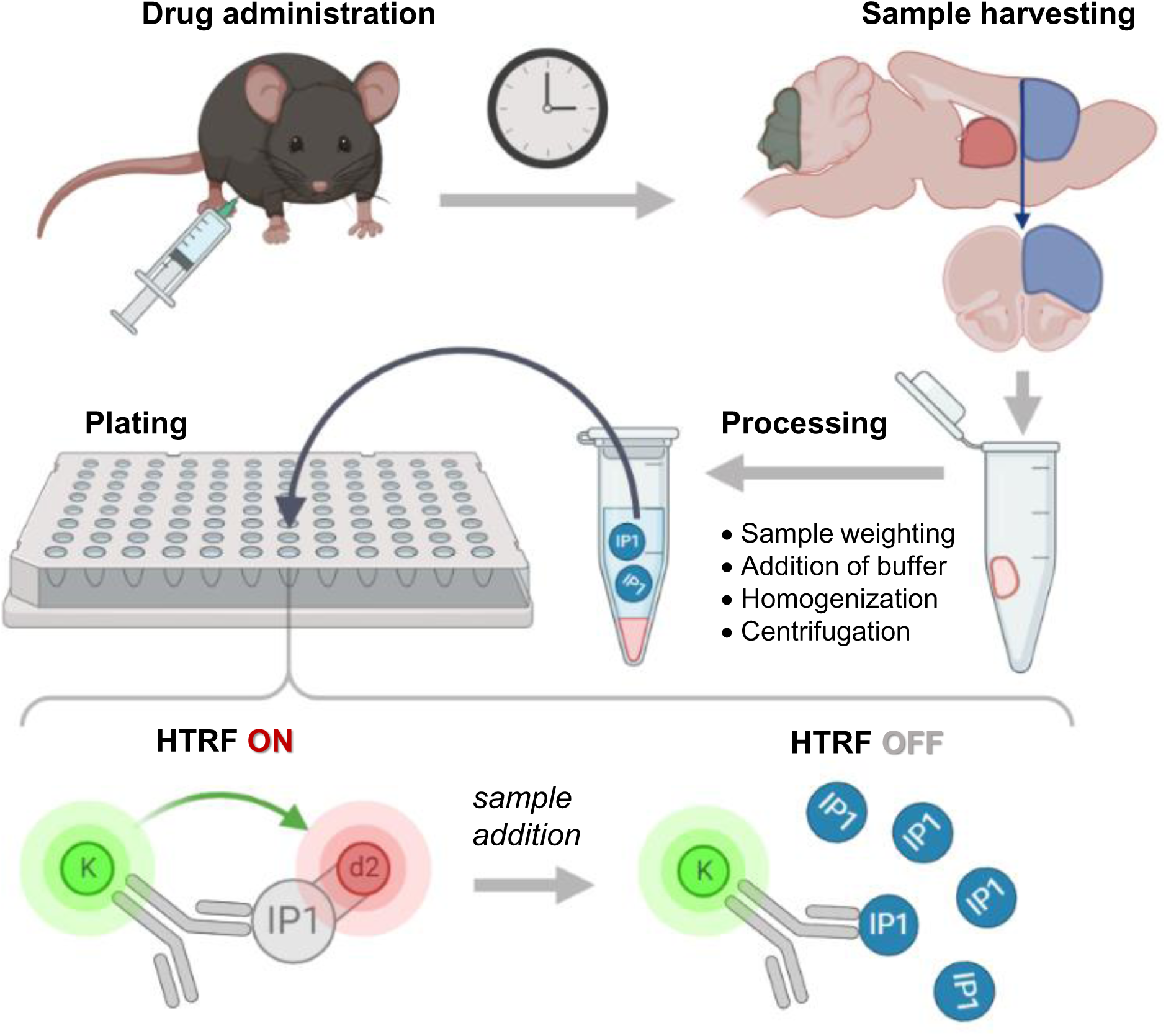
Workflow depicting the methodology. Mice brains are collected and dissected at predefined times after administration of the test drug. The areas dissected corresponding to the frontal cortex (blue), striatum (red) and the cerebellum (green) are shown in the sagittal section. The coronal section indicates the approximate posterior limit of the frontal cortex samples. After processing the sample, an aliquot of the clarified homogenate is transferred to an HTRF plate alongside the corresponding reagents. After incubation, the decrease in HTRF due to the displacement of the acceptor (d2-labelled IP1) from the donor (K-labelled Ab) by biological IP_1_ present in the homogenate is measured in a plate reader. See Methods section for details.

HTRF detection of IP_1_ relies on immunodetection coupled with a fluorescence resonance energy transfer (FRET) system^18^. Two main components integrate this system: a donor Tb-cryptate tagged anti-IP_1_ antibody and an acceptor-tagged IP_1_ molecule. FRET signal is maximal when both components are in aqueous media forming an immunocomplex. When non-labeled IP_1_ is present (*i.e*., from a biological sample), it displaces the acceptor-tagged IP_1_ from the system, and the FRET effect is suppressed (**Fig 1**). By virtue of the long-lived fluorescence emission of a Tb cryptate donor in the system, the assay readout is time-resolved. This millisecond-scale delay in the readout is sufficient to outlast the almost-immediate decay in the fluorescence signal of endogenous biological fluorophores; a common source of interference that precludes the use of fluorescence-based techniques to native tissue preparations^18,19^. The experimental assay window (signal/baseline in standard curve) was >10 (**Supp Fig 1**.).

### Brain region-specificity and 5-HT_2A_R involvement

DOI, or 1-(2,5-dimethoxy-4-iodophenyl)-2-aminopropane, is a potent psychedelic drug and valuable tool in the study of psychedelics’ pharmacology^20^. As a proof-of-concept study in the involvement of 5-HT_2A_R in the action of psychedelics, wild-type (WT) and 5-HT_2A_R knockout (5-HT_2A_R-KO) animals were treated with 2 mg/kg of DOI; a dose known to be active at the molecular and behavioral level, and present in the mouse brain at the time (1h) of sample harvesting ^21,22^.

In the WT cohort, frontal cortex homogenates from DOI-treated animals showed an increased IP_1_ signal as exemplified by the quotient of emission at 615 nm and 655 nm. To further demonstrate specificity of the IP_1_ signal, we compared frontal cortex homogenates between WT and 5-HT_2A_R-KO animals, both treated with DOI (**Fig 2A**). The robust increase in IP_1_ signal in WT frontal cortex homogenates was not replicated in 5-HT_2A_R-KO; which suggests that the main driver of the IP_1_ signal detected in tissue samples from the mouse frontal cortex homogenates is a consequence of 5-HT_2A_R stimulation (two-way ANOVA, genotype: F[1,8]=57.50, P<0.001; treatment: F[1,8]=82.20, P<0.001. Bonferroni’s post hoc (veh vs DOI): WT, P<0.001; 5-HT_2A_R-KO, P>0.05). A similar demonstration of 5-HT_2A_R involvement was previously shown by blockage of IP_1_ signal by the putative psychedelic drug quipazine with pre-treatment with the 5-HT_2A_R antagonist M100907^23^.

**Figure 2.**
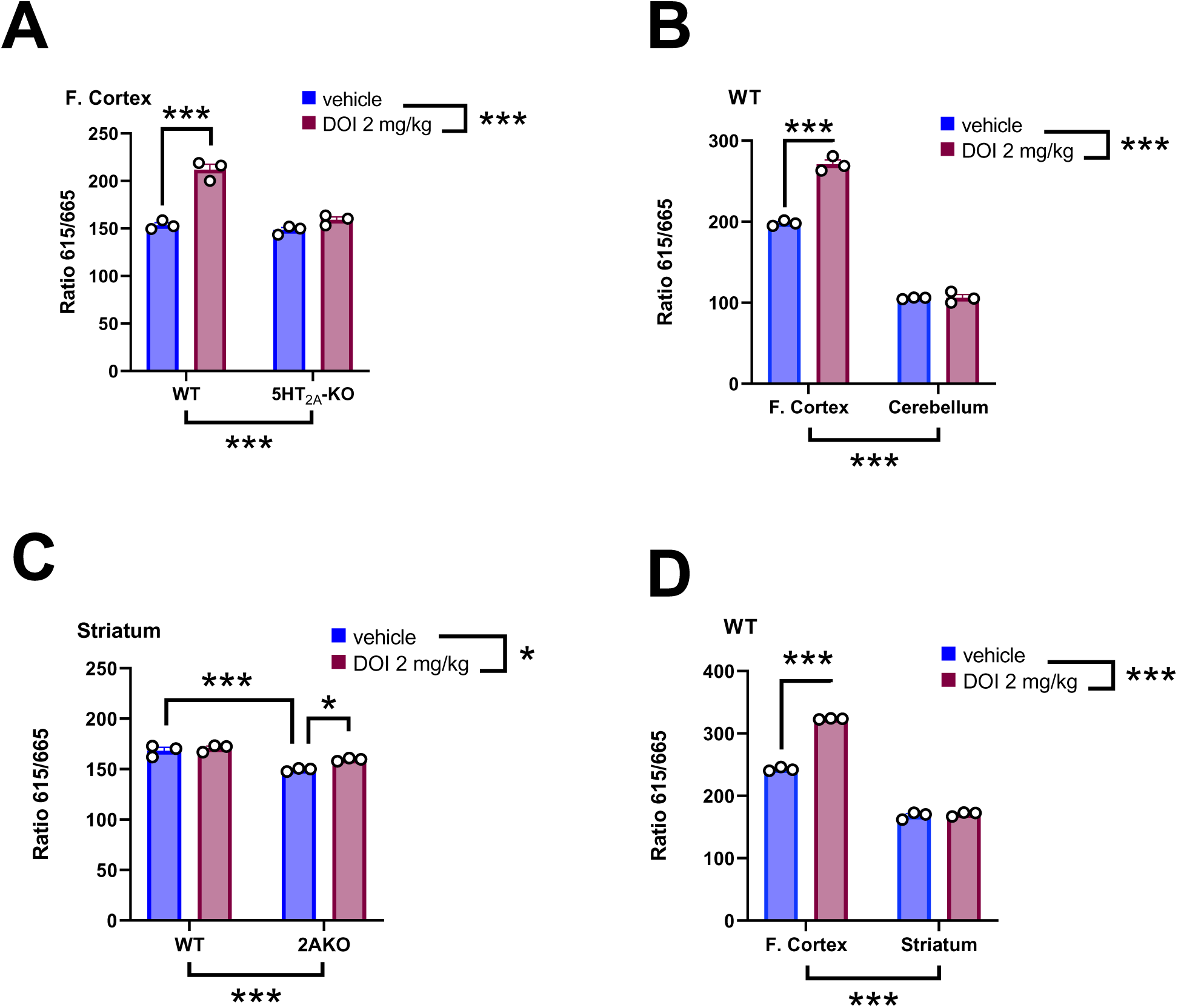
IP_1_ signal from different brain regions harvested from mice (n=3 per group) treated with DOI (2 mg/kg) (i.p) and sacrificed 30 min after drug administration. Differences between WT and 5-HT_2A_R-KO in the frontal cortex (**A**), between the cerebellum and the frontal cortex in WT mice (**B**), between WT and 5-HT_2A_R-KO in the striatum (**C**) and between the frontal cortex and the striatum in WT mice (**D**). Two-way ANOVA with Bonferroni’s post hoc analysis. *P<0.05, ***P<0.001.

These results are also consistent with the dense expression of 5-HT_2A_R-KO in the mouse anterior cortex^24^, and further supported by the absence of changes in the IP_1_ signal in the cerebella of the same animals, where 5-HT_2A_R expression is merely absent (**Fig 2B**) (two-way ANOVA, region: F[1,8]=1421, P<0.001; treatment: F[1,8]=116.0, P<0.001. Bonferroni’s post hoc (veh vs. DOI): F. cortex, P<0.001; Cerebellum, P>0.05). It is also worth noting that baseline levels of IP_1_ are greater in the frontal cortex than in the cerebellum.

We also assessed receptor specificity per brain region by analyzing the striatum. DOI produced a minimal increase in IP_1_ signal in both WT and 5-HT_2A_R-KO animals of comparable magnitude, albeit only significant in 5-HT_2A_R-KOs (**Fig 2C**) (two-way ANOVA, genotype: F[1,8]=54.24, P<0.001; treatment: F[1,8]=9.35, P<0.05. Bonferroni’s post hoc (veh vs DOI): WT, P=0>0.05; 5-HT_2A_R-KO, P<0.05). Interestingly, baseline levels of IP_1_ appeared to be lower in the striatum of 5-HT_2A_R-KO mice compared to WT controls. Responsiveness to DOI and baseline levels greatly differed between the frontal cortices and striata of WT mice (**Fig 2D**) (two-way ANOVA, region: F[1,8]=2785, P<0.001; treatment: F[1,8]=374.5, P<0.001. Bonferroni’s post hoc (veh vs DOI): F. cortex, P<0.001; Striatum, P>0.05). In contrast to the cortex, 5-HT_2A_R expression is much less abundant in the striatum^24^, a region known for the abundance of 5-HT_2C_Rs^25^, another G_q_-coupled serotonin receptor activated by psychedelics, including DOI.

### Time-course of IP_1_ in the mouse brain

As a downstream readout in vivo, we expected that the IP_1_ signal would be influenced by the pharmacokinetic constraints that modulate the action of the stimulating drug^21^. To gain further insights on the temporal dynamics of DOI on this readout, we evaluated the evolution of IP_1_ signal over time in mice administered with 2 mg/kg of DOI (**Fig 3**). IP_1_ signal increased sharply during the first 1h post administration to then progressively decrease and return to baseline levels at >6h). Compared to the characteristic biphasic pharmacokinetic curve^21^, the time-course of IP_1_ in frontal cortex homogenates appeared to show truncated peak of effect (*i.e*., flattened maximum). This could indicate saturation of the biological system having reached the maximal IP_1_ production possible, rather than saturation of the signal (see **Supp Fig 1**. as an example of assay window).

**Figure 3.**
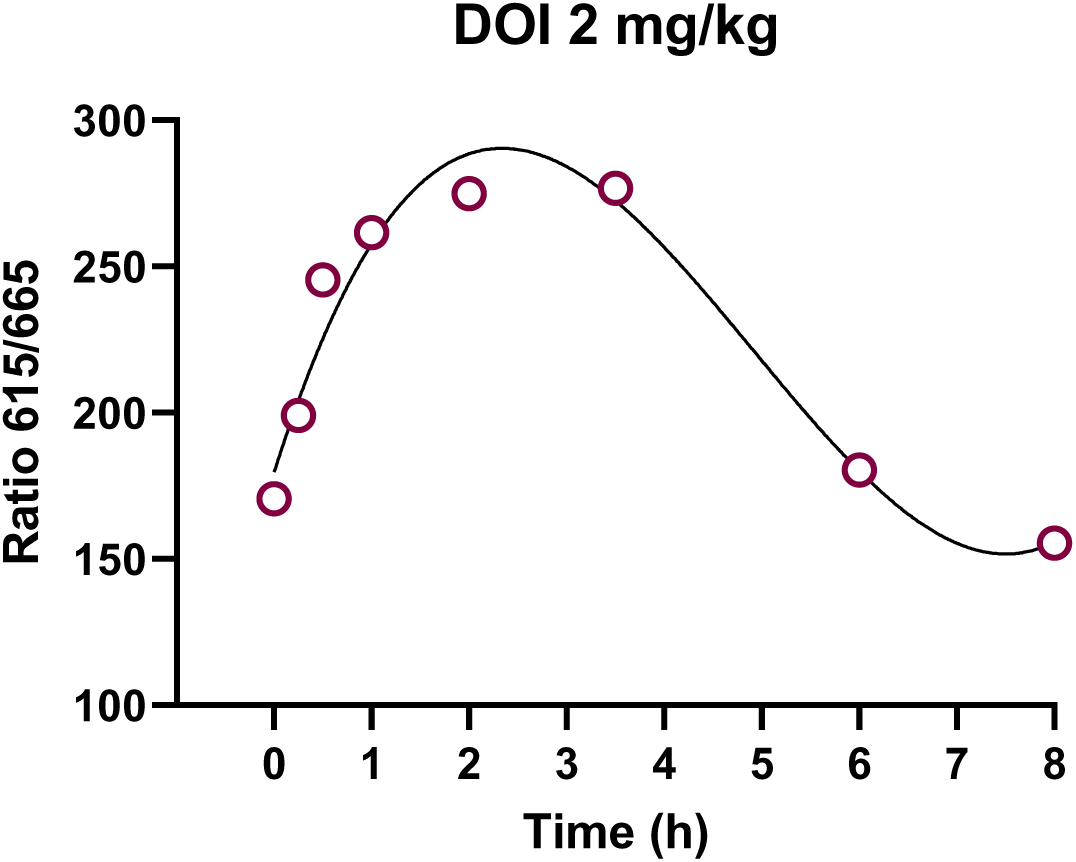
Time-course of IP_1_ signal from frontal cortices of WT treated with DOI (2 mg/kg) (i.p) and sacrificed at different times after drug administration (n=3 technical replicates, one animal per timepoint). Non-linear fitted third order polynomial curve shown.

### Dose-response evaluation and behavioral correlates

DOI robustly increases HTR in mice and is a potent 5-HT_2A_R agonist in heterologous expression models in vitro^21^. To further explore the molecular and behavioral link of DOI 5-HT_2A_R action, we sought to determine HTR correlativeness to IP_1_ signal induced by different doses of DOI within the same subjects (**Fig 4A**).

**Figure 4.**
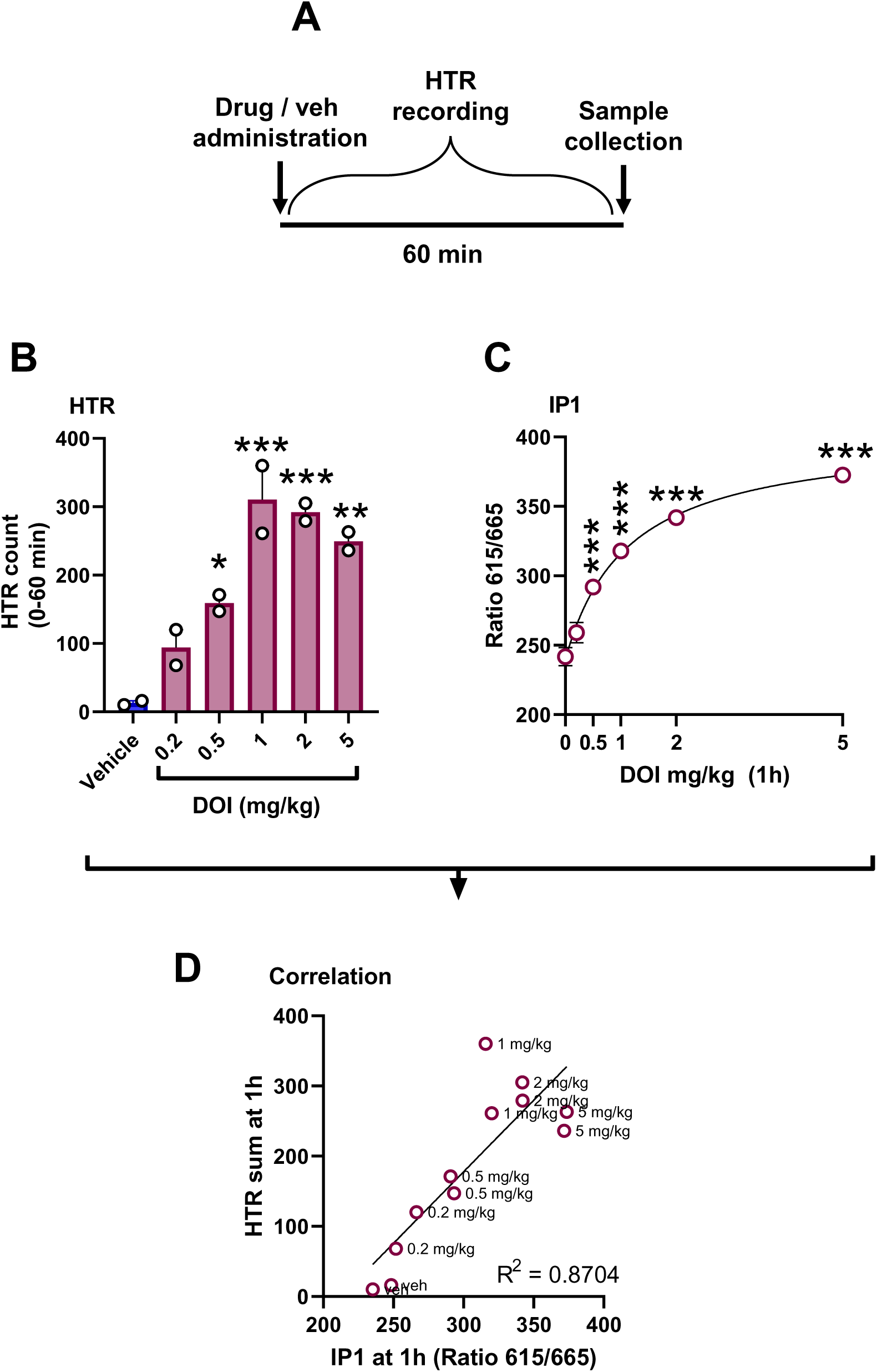
Scheme of the experimental design to determine HTR and IP_1_ in the frontal cortex of the same animals (n = 2 per dose, WT) 1h after DOI administration (**A**). Cumulative HTR counts (**B**). IP_1_ signal in the frontal cortices and fitted saturation curve (**C**). Correlation analysis of cumulative HTR counts and IP_1_ in the frontal cortex per animal after 1h since DOI administration (**D**). One-way ANOVA with Bonferroni’s post hoc analysis vs vehicle. *P<0.05, **P<0.01, ***P<0.001.

As expected, increasing doses of DOI resulted in higher HTR counts; as determined by their electromagnetic signature. HTR plateau was reached at ∼1 mg/kg after 60 min (**Fig 4B**) (**Supp Fig 2**) (one-way ANOVA, F[5,6]=23.10, P<0.001. Bonferroni’s post hoc (veh vs DOI): 0.2 mg/kg, P>0.05; 0.5 mg/kg, P>0.05; 1 mg/kg P<0.001; 2 mg/kg, P<0.001; 5 mg/kg, P<0.01).

Immediately after recording HTR for 60 min after administration of DOI, the animals were sacrificed and the frontal cortices harvested. IP_1_ signal at 60 min followed an asymptotic dose-response pattern (**Fig 4C**) (one-way ANOVA, F[5,6]=142.3. Bonferroni’s post hoc (veh vs DOI): 0.2 mg/kg, P=0.08; 0.5 mg/kg, P<0.001; 1 mg/kg P<0.001; 2 mg/kg, P<0.001; 5 mg/kg, P<0.001). Furthermore, IP_1_ signal readout was highly correlated with the 60 min aggregated HTR counts (**Fig 4D**)(F[1,10]=23.91, P<0.001).

### Evaluation of other serotonergic drugs

LSD is among the most paradigmatic examples of classical psychedelics. Its core ergoline structure is shared by lisuride. Lisuride, despite being a non-psychedelic analogue, shares with LSD its ability to potently bind to and activate the 5-HT_2A_R in vitro^6,26^. We then sought to evaluate this apparent paradox in the mechanism of action of psychedelics by determining in parallel IP_1_ signal in the mouse frontal cortex for both drugs at different doses (**Fig 5A**) (two-way ANOVA, dose: F[3,8]=103.7, P<0.001, treatment: F[3,8]=442.9. Bonferroni’s post hoc (veh vs LSD): 0.2 mg/kg, P<0.001; 0.4 mg/kg, P<0.001; 1 mg/kg, P<0.001. Bonferroni’s post hoc (veh vs lisuride): 0.2 mg/kg, P<0.05; 0.4 mg/kg, P=0.0397; 1 mg/kg, P<0.001). Fitting IP_1_ signal to a non-linear regression model showed a ∼6-fold differences in the saturation magnitude between both drugs (LSD, ED_50_=0.1191 mg/kg, span=58.28 ratiometric arbitrary units (A.U.), R-square=0.9837; lisuride, ED_50_=0.3401 mg/kg, span=20.82 A.U., R-square=0.8475). The meager production of IP_1_ signal by lisuride contrast with the robust increase shown by LSD.

**Figure 5.**
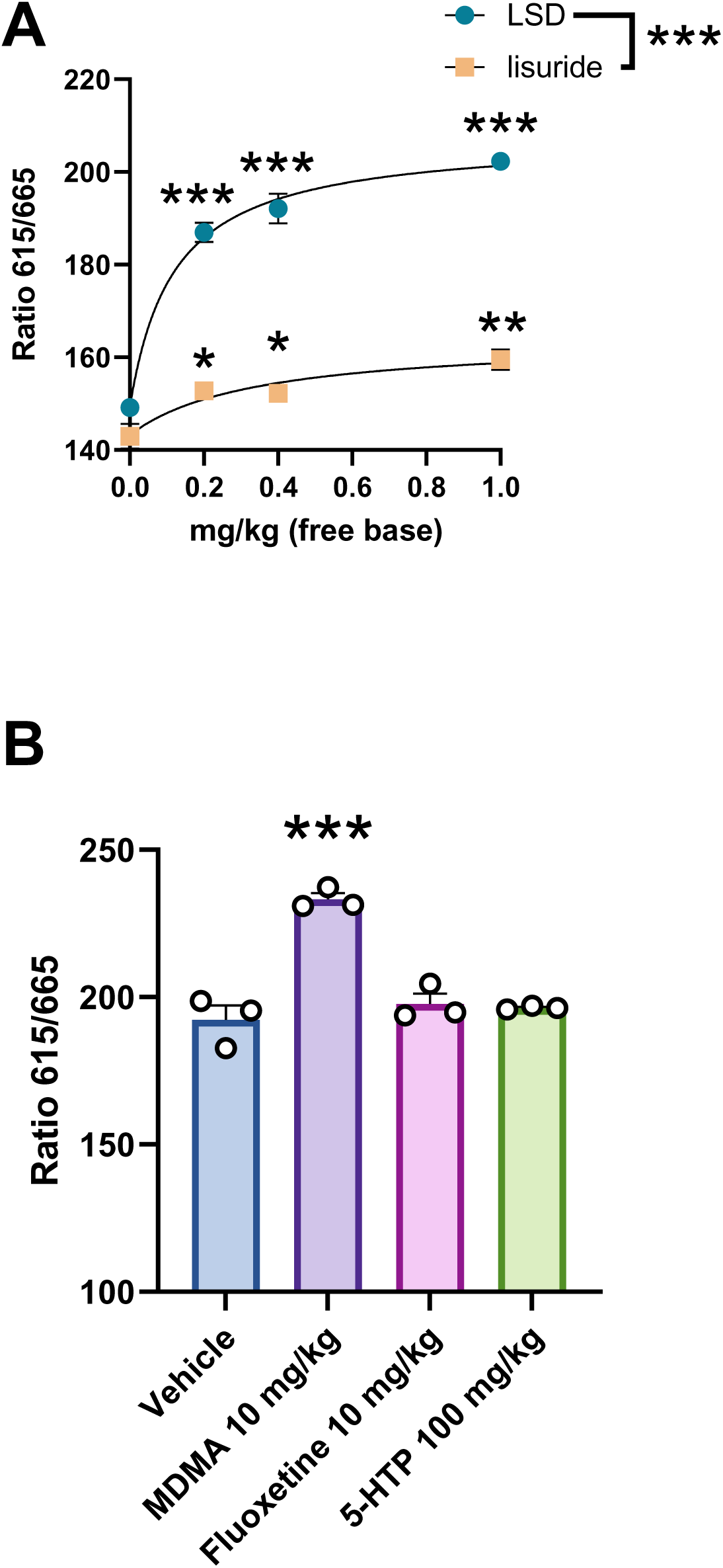
IP_1_ signal 1h after drug administration from the frontal cortex of mice (n=2 per dose) treated with different doses of LSD and lisuride (**A**) and fitted saturation curve. IP_1_ signal 1h after administration of different serotonergic drugs (n=3 per drug group) (**B**). Two-way (**A**) and one-way (**B**) ANOVA with Bonferroni’s post hoc analysis vs vehicle. *P<0.05,*** P<0.001.

To further explore the versatility of the platform, we sought to evaluate the IP_1_ signal for different serotonergic drugs neighboring the classical psychedelics chemical space (**Fig 5B**): MDMA, fluoxetine and 5-hydroxytryptophan (5-HTP). MDMA is a serotonin releaser that reverses serotonin transport thus leading to increases in the neurotransmitter in the synaptic space^27^. The traditional antidepressant fluoxetine directly blocks the serotonin transport back to the synapse^28^. 5-hydroxytryptophan (5-HTP) is metabolic intermediate in the biosynthesis of serotonin. Despite its lack of psychedelic effects in human at doses commonly used as a dietary supplement, 5-HTP induces HTR in mice at high doses, thus constituting one paradigmatic example of false positives in this predictor model of psychedelic effects in human^29,30^.

Out of the different drugs tested, only MDMA produced a substantial increase in IP_1_ signal (**Fig 5B**) (one-way ANOVA, F[3.8]=36.08, P<0.001. Bonferroni’s post hoc (vehicle vs drug): MDMA, P<0.001, fluoxetine, P>0.05; 5-HTP, P>0.05). The matched dose of fluoxetine, a dose known to produce behavioral effects in mice^31^, did not produce any apparent changes in the levels of IP_1_. Surprisingly, neither did 5-HTP at a dose known to induce HTR increases and intense diarrhea in mice^29^.

To ensure proper coverage of 5-HTP pharmacokinetics, in a separate experiment, 5-HTP was administered to WT mice at a higher dose (200 mg/kg) and samples collected at an earlier time-point (30 min) following recording of HTR (**Fig 6A**). 5-HTP produced a sharp increase of HTR events compared to vehicle, as expected (**Fig 6B**) (**Supp. Fig 3**)(P<0.01, t=4.656, df=4). However, the HTR induction profile was not paralleled by an increase in IP_1_ signal in the frontal cortex homogenates (**Fig 6C**)(one-way ANOVA, F[2,5]=87.21, P<0.001; Bonferroni’s post hoc (vehicle vs DOI), P<0.001).

**Figure 6.**
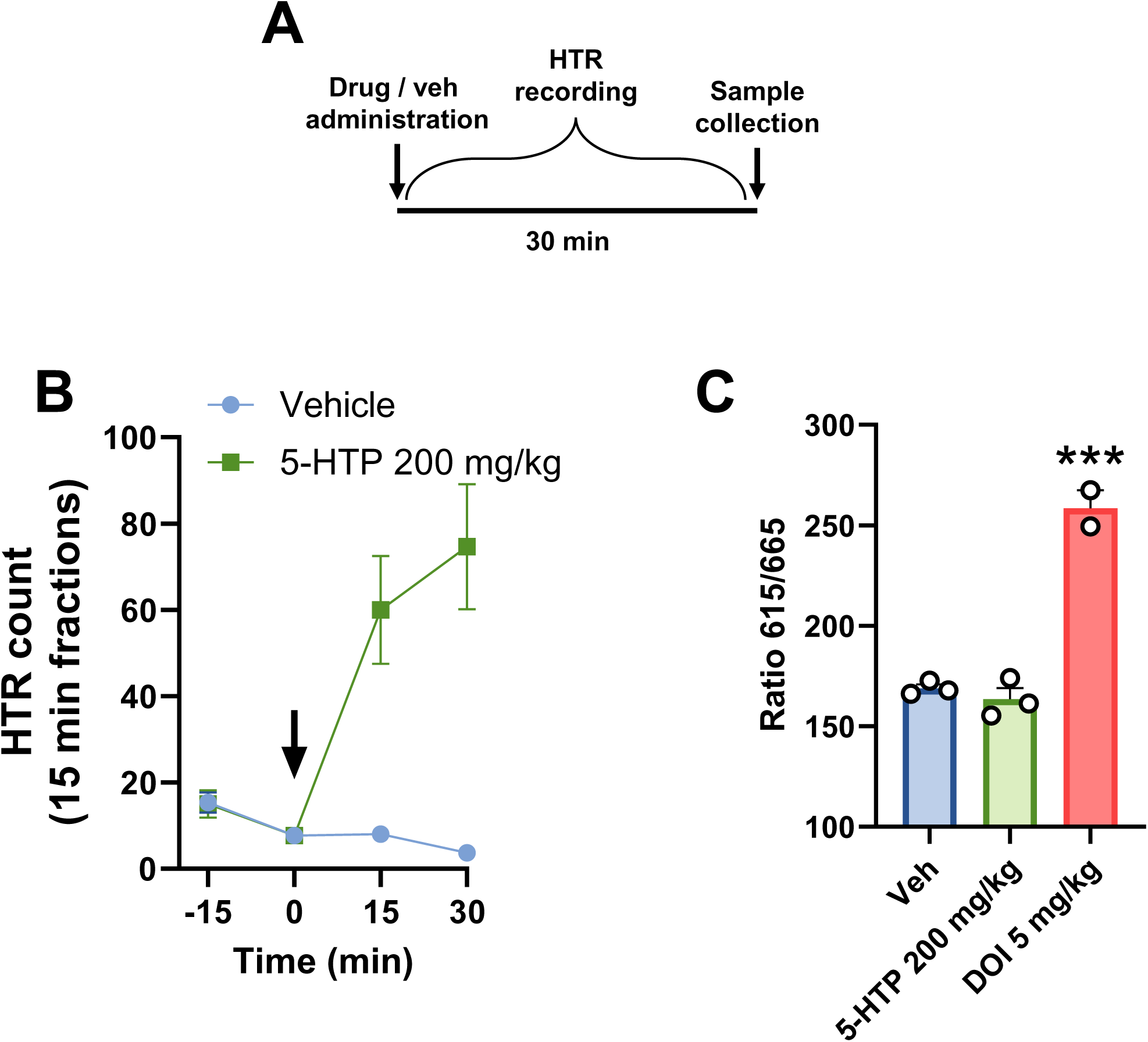
Scheme of the experimental design (**A**) to quantitate HTR (**B**) and IP_1_ (**C**) in the frontal cortex of the same animals 30 min after the administration 5-HTP (200 mg/kg) (n = 3). DOI 5 mg/kg (n = 2) as positive control (**C**). One-way ANOVA with Bonferroni’s post hoc analysis vs. vehicle. **P<0.01.

## Discussion

Activation of 5-HT_2A_R by classical serotonergic psychedelic drugs is the necessary pharmacological driver for their distinct effects in human psyche^5,32^. Canonically, 5-HT_2A_R activation results in G_q_ protein isoforms coupling to the receptor triggering a cascade of downstream events that culminates with the hydrolysis of IP_1_ ^6,33^. Herein, we present a methodology that enables capturing a snapshot of IP_1_ levels in mouse brain homogenates as a surrogate readout of 5-HT_2A_R activation in mouse cortex. As a proof of concept, we demonstrate that increases in IP_1_ can readily be detected in frontal cortex brain homogenates following administration of psychedelics LSD and DOI to the alive animal. We also demonstrate the involvement of 5-HT_2A_R in this readout for DOI by genetic deletion of the receptor and by differential expression of the receptor in different neuroanatomical regions. DOI produced a distinct dose-dependent increase of IP_1_ in the 5-HT_2A_R-dense frontal mouse cortex in comparison with the cerebellum, a region known to lack in expression of 5-HT_2A_R^25^. The receptor specificity shown is in alignment with our previous study showing that IP_1_ signal in frontal cortex of mice treated with the 5-HT_2A_R agonist quipazine was blocked by pre-treatment with the selective antagonist M100907^23^.

Our results demonstrate that IP_1_ measured by HTRF in cortical homogenates can afford sufficient receptor and anatomical specificity to be utilized as a potential biomarker of 5-HT_2A_R activation in the frontal cortex. This prospective use is not without limitations. Psychedelics target several other GPCRs expressed in the central nervous system that upon activation couple G_q_ proteins. These considerations are important when considering potential experimental designs. We observed a small but statistically significant effect of DOI on IP_1_ signal in the striatum of 5-HT_2A_R-KO that could be attributed to activation of the 5-HT_2C_R. As it pertains to the use of frontal cortex as a region of interest to characterize the pharmacology of 5-HT_2A_R agonists in general, and psychedelics in particular, 5-HT_2C_R did not appear to contribute to the frontal cortex turnover of IP_1_ as experimentally shown with DOI in WT and 5-HT_2A_R-KO mice. This is likely possible due to the relative abundance of 5-HT_2A_R in the cortex relative to 5-HT_2C_R expression^25,34^.

We encountered that HTR and IP_1_ in frontal cortex homogenates from the same mice treated with DOI were highly correlated at 1h post administration. We appreciated divergences in the kinetics, however. Consistent with the inverted U-shape of HTR dose responses^35^, HTR counts appeared to drop at higher doses of DOI. Conversely, the IP_1_ signal in the frontal cortex of the animals followed an asymptotic fate akin to a saturation curve. It is plausible that behavioral disruption limits the expression of HTR at doses that might continue to stimulate 5-HT_2A_R before reaching saturation of the biological system. In the time domain, IP_1_ turnover increase was delayed relative to manifestation of HTR and our previously reported DOI brain pharmacokinetics^21^. IP_1_ is a near-terminal metabolite of the G_q_ pathway so it is possible that its formation can be subject to downstream catabolic bottlenecks. As we recently reported, exposure to DOI leads to downregulation of 5-HT_2A_Rs in the frontal cortex and development of HTR tolerance^36^. IP_1_ formation might thus continue intracellularly after clearance of the drug-bound receptor from the cell surface.

Furthering in the study of psychedelic effect correlative to 5-HT_2A_R activation, we also tested LSD and lisuride in our IP_1_ model. Both compounds are brain penetrant ergolines and 5-HT_2A_R agonists, yet LSD is one of the most paradigmatic representatives of the psychedelic drug family, whereas lisuride is not psychedelic in human or animal models^6,37^. Increasing doses of LSD produced a saturable IP_1_ signal in the 5-HT_2A_R-rich mouse frontal cortex that was not matched in magnitude by lisuride. The lesser ability of lisuride to activate 5-HT_2A_R receptors in vivo might ultimately answer its lack of LSD-like effects as well as its apparent 5-HT_2A_R antagonist profile in numerous paradigms such as spontaneous HTR^6,36^.

We also screened several other serotonergic compounds and measured their effect on IP_1_ in mouse frontal cortex homogenates following systemic administration. Only MDMA at 10 mg/kg produced a statistically significant increase in IP_1_ compared to vehicle treated animals. Both MDMA and fluoxetine target the serotonin transporter primarily but differ in their primary effect on serotonin’s homeostasis. Fluoxetine blocks its uptake^38^ whereas MDMA produces serotonin efflux from its presynaptic storage^39^. It is plausible that the latter mechanism could contribute to greater levels of 5HT capable of attaining meaningful stimulation of cell signaling via 5-HT_2A_R in the mouse frontal cortex. Unlike fluoxetine, MDMA shares certain aspects of its phenomenological experience with classic psychedelics^40^ some of which can be blocked by the 5-HT_2_R antagonist ketanserin^41^.

HTR is one of the best classifiers of psychedelic effect in human for serotonergic drugs, but it is not exempt from the interference from false positives^35^. One of such cases is 5-HTP^29,30^. Devoid of psychedelic effect in human at doses used as a ‘nutritional supplement’, 5-HTP induces in mice HTR sensitive to the action of 5-HT_2A_R antagonists^30^. However, we showed that a HTR-inducing dose of 5-HTP did not stimulate 5-HT_2A_R-dependent IP_1_ signaling in the mouse frontal cortex. Our findings suggests that 5-HTP-induced HTR could be extra-cortical in nature. This also demonstrates how IP_1_ detection in mouse brain samples can complement the predictive power of HTR in the determination of psychedelic potential in human.

The study of 5-HT_2A_R pharmacodynamics at the receptor level generally starts with the use of in vitro approaches to avoid confounders in the readout. Singling out the receptor in heterologous expression in these models detach functional readouts from its physiological context. For identical compounds, historical divergences in reported maximal efficacies are common even for readouts that belong in the same G_q_-dependent signaling pathway and heterologous expression system. One fundamental advantage of our model is that it provides a representative snapshot of IP_1_ levels in the brain. The readout integrates a wide array of intervening neurobiological and pharmacokinetic processes that occur in the entire animal that are not accounted for in cell systems. While animal intensive, the method does not require any additional experimental interventions besides drug administration during the in-life period and sample harvesting, thus avoiding introducing secondary sources of variability that can operate as external confounders. Other ex vivo techniques employed in the quantitation of inositol phosphates in the rodent brain involve incubation with the test compound in an organ bath^42^. In our case, sample processing is limited to homogenization following brain extraction and dissection to ensure that IP_1_ levels are representative of the drug effect during the in-life period. It also does not require the use of radioactive reagents and affords a readout of IP_1_ levels in less than 1h from sample collection.

The methodology is not exempt from some limitations, that while addressable, are worth considering. For instance, IP_1_ quantification alone provides no hints on the stoichiometry or receptor and G_q_ protein isoforms responsible for the readout. In vitro approaches using heterologous expression systems are better suited to study such a degree of molecular precision ^43^. Another potential caveat in our approach is that 5-HT_2A_R is not the only receptor capable of coupling G_q_ proteins in the frontal cortex with the consequential increase in IP metabolites. This calls for a careful evaluation of the drug-selectivity profile, neuroanatomical expression of target and off-target receptors and ultimately the test drug pharmacokinetics^17^. However, as exemplified here with DOI, these confounders can be controlled through experimental design in a way that can also produce valuable mechanistic insights. Quantitation of IP_1_ in the mouse brain, as shown here, can complement in vitro approaches and expand our understanding of 5-HT_2A_R pharmacodynamics for psychedelics and related compounds.

### Conclusions

As psychedelic research gains traction, the need for complementary models that can afford a better understanding of their pharmacology becomes increasingly apparent. In response, we developed a platform capable of capturing a snapshot of drug mediated. 5-HT_2A_R activation of the G_q_ pathway in the mouse cortex via detection of IP_1_ in tissue homogenates. This methodology aims to bridge the gap between 5-HT_2A_R in vitro and in vivo pharmacodynamics by using a model that employs a molecular readout in the mouse brain that subjects the drug to the same pharmacokinetic constraints that drive behavioral pharmacology. We demonstrated a high degree of 5-HT_2A_R specificity in DOI stimulation of IP_1_ in the mouse frontal cortex that enables the exploration of this readout as a reference for the quintessential molecular interaction driving the subjective effects of psychedelics. It also serves as a comparator for other serotonergic drugs with related mechanisms of action. The platform can aid the exploration of relationships between drug-mediated 5-HT_2A_R activation, pharmacokinetics and dynamics of behavioral manifestations in the same animals.

## Methods

### Animals

Wild-type C57BL/6J male mice and 5-HT_2A_R-KO ^6^ male mice backcrossed from 129S6/SvEv onto C57BL/6J for several generations (F5) were randomly allocated into the different treatment groups (12-16 weeks old). Animals were housed on a 12 h light/dark cycle at 23°C with food and water ad libitum. All procedures were conducted in accordance with NIH guidelines and were approved by the Virginia Commonwealth University Animal Care and Use Committee. All efforts were made to minimize animal suffering, and the number of animals used.

### Drugs

Drugs were administered dissolved in 0.9% saline as vehicle intraperitoneally (5 ul/g, i.p.) and sourced from authorized vendors: (±)-1-(2,5-dimethoxy-4-iodophenyl)-2-aminopropane ((±-DOI) hydrochloride (MilliporeSigma); LSD (Lipomed); lisuride maleate (Tocris); 5-HTP (MilliporeSigma); fluoxetine hydrochloride (Tocris); (±)-MDMA hydrochloride (Lipomed). An equimolecular amount of HCl was added to form the hydrochloride salt in situ for LSD and 5-HTP. In the case of LSD and lisuride, the vehicle contained an equivalent volume of DMSO (<1% final volume).

### Quantification of head-twitch responses (HTR)

HTR assessment was performed in mice with magnetic ear tags designed for automated HTR detection^21,44^. Briefly, neodymium magnets bearing-ear tags (∼50 mg) were placed bilaterally through the pinna antihelix under isoflurane (2%)^44^. All animals were allowed to recover for a week prior to testing^21^. During the testing session, animals were transferred from their home cage to the testing chamber where they were allowed to habituate to the environment. After 30 min, DOI or 5-HTP or vehicle was administered i.p.. Data acquisition in the magnetometer was performed for 60 min for DOI and 30 min for 5-HTP as previously described^21^. After completion of the recording, the animals were sacrificed by cervical dislocation, and the brain samples harvested. Data was processed offline using a previously described signal analysis protocol^44^. To refine HTR detection, the signal was also processed using a deep learning-based protocol based on scalograms^45^. Mismatches between both detection methods were inspected visually without clues relative to the timestamp of the event or treatment group as previously described^21,44^.

### IP1 detection in brain sample homogenates

#### Sample collection

Animals treated with test drug or vehicle were sacrificed at the time specified on each experimental design by cervical dislocation. All animals in each experiment and time point were sacrificed simultaneously (∼3 animals/min), decapitated, and the heads cooled down on wet ice. The brain extraction and dissection of the region/s of interest (**Fig 1**.) was performed sequentially. During this process, only saline 0.9% was employed for washing of specimens as the use of PBS can influence the HTRF readout. For consistency, during processing of a batch, samples and heads to be processed remained on ice until the dissection process was completed, and then samples were frozen at - 80°C for storage. The carryover of saline, blood or any other debris was minimized during this process. Quality and homogeneity in the dissection process proved crucial to obtain accurate readouts.

#### Sample processing

Each individual sample was transferred frozen to a 1.5 ml conical tube with a safelock cap (#3456 ThermoFisher) in a semi-micro balance (MS105DU, Mettler Toledo) to weight the sample with precision. A volume of 0.5 mm diameter glass beads (Biospec products #11079105) roughly equivalent to the sample volume and an exact amount of chilled homogenization buffer (10 μl/mg of sample) was added to the same tube. Homogenization buffer was prepared as a mix of 10% ‘Lysis & Detection Buffer’ and 90% ‘Stimulation Buffer’ 1× from the IP-One Gq kit (Perkin Elmer). While the exact formula of this buffers is proprietary, they contain LiCl to prevent IP_1_ enzymatic hydrolysis as outlined in the user manual. The precision in the volume-to-weight dilution proved to be crucial for consistency and narrowing of standard error of the mean. The tubes caps were secured and the samples homogenized (NextAdvance Bullet Blender 24) at 4C for 5 min, speed 6, after which they were centrifuged for 15 min microcentrifuge (4C, 17,000 g). The clarified homogenate supernatant was used directly or frozen at −80C until further use.

#### Platting and reading

The detection reagents: donor Tb cryptate antibody (K) and d2-labelled acceptor (d2) were reconstituted according to the IP-One Gq kit manufacturer instructions in distilled water (6X) and then diluted in ‘Lysis & Detection Buffer’ (1X). Aliquots were kept frozen and 20C. Platting was performed in white opaque HTRF 96 well low volume plates (66PL96005). Each well contained 18 μl of a mastermix composed by 12 μl of ‘Stimulation Buffer’ 1×, 3 μl of K reagent, 3 μl of d2 reagent. To reach the 20 μl final volume per plate, 2 μl of clarified homogenate supernatant (sample) or an aliquot of known concentration of IP_1_ (standard) were added. Mastermix was prepared fresh before each experiment and standard curves generated (See **Supp.** Fig. 1) Each experiment had, at least, a 0 μM and a 22 μM IP_1_ standard as internal control. The plate was covered and incubated for 30 min in darkness at room temperature and read in a VICTOR Nivo plate reader (Perkin Elmer). The incubation is not involved in the production of IP_1_, addition of DOI 10 μM prior to incubation to control wells did not change IP_1_ levels (results not shown). The settings employed for the reading were: excitation filter 320/8 nm, emission filter #1 615/8 nm, emission filter #2 665/8 nm, dichroic mirror D400, delay time 70 μs, emission time 200 μs, flash energy low, measurement time 500 ms, two reads per well, Z-focus 12 mm. The ratiometric signal was calculated for each well as the reading from Emission filter #1 divided by the reading from Emission filter #2 and multiplied by 100.

#### Statistical analysis

Statistical significance involving three or more treatments or different doses of the same treatment was assessed by one-way ANOVA followed by Bonferroni’s post hoc test. Statistical significance of experiments involving different treatments and doses was assessed by two-way ANOVA followed by Bonferroni’s post hoc test. The level of significance was set at p = 0.05. All values in the figure legends represent mean ± s.e.m. Statistical analysis was performed with GraphPad Prism software version 9 (La Jolla, CA).

## Author contributions

M.F.R. conceived the methodology, performed the experiments, analyzed the data and wrote the manuscript along with J.G.-M. who supervised the research and obtained funding.

## Acknowledgements

This work was supported by the National Institute of Health (NIH) grants T32MH020030 (M.F.R.) and R01MH084894 (J.G.-M.).

---

Supp Fig 1. Standard curve corresponding to different concentrations of IP_1_ from a standard used as internal controls from three different experiments. Shown are the corresponding non-linear fit curves and parameters.

Supp Fig 2. Time-course of vehicle and DOI-induced HTR at different doses. Arrow shows the point at which the drug is administered.

Supp Fig 3. Sum of HTR events for 30 min post-administration of 5-HTP (200 mg/kg) or vehicle. **P<0.01.

