## Supplemental Figures for "Snapshot of 5-HT_2A_ receptor activation in the mouse brain via IP_1_ detection"

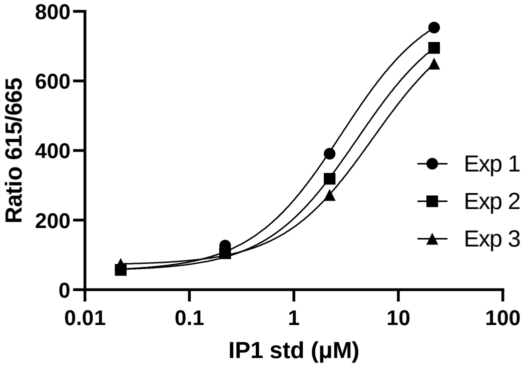


| | Exp 1 | Exp 2 | Exp 3 |
|---|---|---|---|
| Bottom | 53.46 | 54.98 | 71.66 |
| Top | 844.2 | 812.1 | 799.4 |
| EC50 | 2.876 | 4.052 | 5.721 |
| logEC50 | 0.4587 | 0.6076 | 0.7574 |
| Span | 790.8 | 757.2 | 727.8 |

**Supp. fig 1.** de la Fuente Revenga et al.

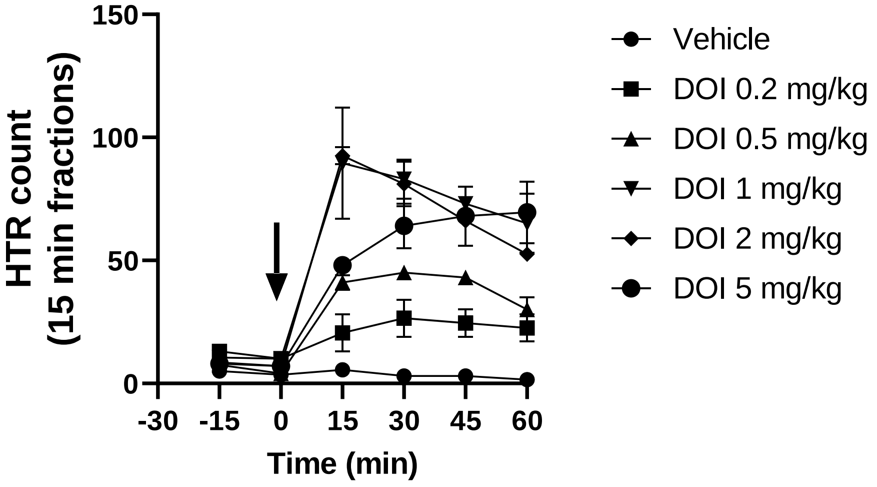


**Supp. fig 2.** de la Fuente Revenga et al.

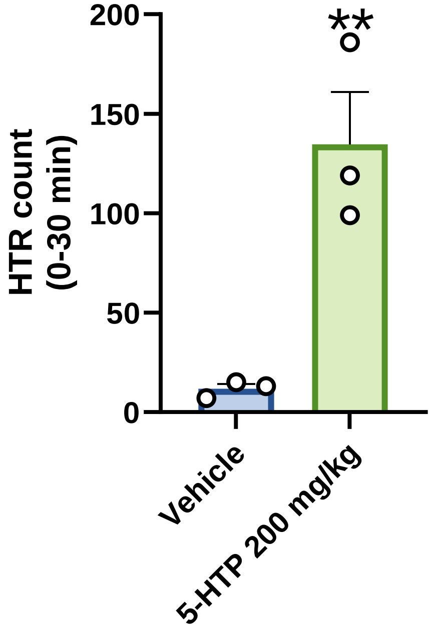


**Supp. fig 3.** de la Fuente Revenga et al.
